# SuperStructure: a Parameter-Free Super-Structure Identifier for SMLM Data

**DOI:** 10.1101/2020.08.19.254540

**Authors:** M. Marenda, E. Lazarova, S. van de Linde, N. Gilbert, D. Michieletto

## Abstract

Single-Molecule Localisation Microscopy (SMLM) allows the quantitative mapping of molecules at high resolution. However, understanding the non-random interaction of proteins requires the identification of more complex patterns than those typified by standard clustering tools. Here we introduce SuperStructure, a parameter-free algorithm to quantify structures made of inter-connected clusters, such as protein gels. SuperStructure works without a priori assumptions and is thus an ideal methodology for standardised analysis of SMLM data.

---

Single-molecule localisation microscopy [1–3] is now commonly employed for quantitative analysis of molecular interactions both in vivo [4–6] and in vitro [7, 8]. SMLM achieves resolutions far beyond the diffraction limit and its typical output is a list of 3D coordinates (or localisation events) that are naturally analysed using efficient clustering algorithms borrowed from quantitative big-data analysis and even astronomy [9–14]. However, clustering algorithms rely on user-defined parameters intrinsically intertwined with the notion of similarity used to define clusters. These parameters can be either assumed or inferred via pre-emptive analysis, yet their choice has a significant impact on the results. This hinders the portability and ease of comparison of different datasets.

At the same time, recent evidence suggest that assemblies of proteins [6, 15–20] and chromatin [21, 22] form complex structures that cannot be simply analysed with standard clustering algorithms. For example, the hnRNP protein SAF-A is suggested to form a dynamic mesh-like structure that maintains transcriptionally active genomic loci in a decompacted configuration [23, 24] whilst superresolution techniques indicates that chromatin is organised in connected nano-scale compartments [25–27]. To understand the relationship between these structures and the dynamics and function of the genome [28–30], a more sophisticated and standardised analysis of SMLM data is required.

Motivated this, we have developed SuperStructure. This method extends the popular density-based clustering algorithm DBSCAN implementing (i) a parameter-free detection and quantification of super-structures made of connected clusters in SMLM data, (ii) a parameter-free quantification of the density of molecules within clusters. Here we show in detail that SuperStructure can be applied to discern and compare complex structures, such as nuclear networks or gels, emerging from different nuclear proteins or cellular conditions, and suggest it is well suited as tool for standardised SMLM analysis.

The working principle of SuperStructure is best explained as follows: while the classic DBSCAN algorithm detects clusters for a fixed choice of minimum number of neighbours *N*_min_ and neighbourhood distance *ϵ* [31], we argue that there is previously overlooked information in how the number of detected clusters *N*_*c*_ changes with *ϵ* for a fixed *N*_min_. By setting *N*_min_ = 0, *N*_*c*_(*ϵ*) is necessarily a monotonically decreasing function as for *ϵ* = 0 every localisation is a cluster by itself and increasing *ϵ* yields fewer but larger clusters. Importantly, the rate at which *N*_*c*_ decays with *E* is an indicator of how quickly localisations, and then clusters of localisations, coalesce with each other and thus of how “connected” they are.

The decay curves outputted by SuperStructure identify different clustering regimes: (i) merging localisations within clusters (intra-cluster regime); (ii) coalescing clusters into super-structures (first super-cluster regime); (iii) merging super-clusters into higher-order super-structures (second/third super-cluster regimes). The rate of decay of *N*_*c*_ in regime (i) is intimately related to the density of emitters *ρ*_*em*_ within the clusters (see Supplementary Note 2), while the decay in regimes (ii) and (iii) are highly dependent on the connectivity between (super-)clusters, as well as on the density of (super-)clusters (see Supplementary Note 3). These two contributions can be further discerned, as we shall see below. See Supplementary Note 1 for details of the algorithm and the pipeline.

We first evaluated the performance of SuperStructure on artificial datasets consisting of inter-connected clusters of simulated localisations (see also Supplementary Note 3). The main dataset is made by connected and spatially homogeneous clusters that are randomly positioned with a density to *ρ*_*cl*_ = 8.2*µ*m^*−*2^ and a radius *R*_*c*_ *∼* 40 *nm*. Pairs of clusters were connected with probability *p* by a sparse points distribution and only if the distance between the clusters was less than *b* = 1 *µm*. This allowed us to readily tune the degree of “connectivity” in the system by varying a single parameter *p* (see also Supplementary Note 3 for another example). The length-scale associated to density of emitters inside clusters *ρ*_*em*_ and to the connections *ρ*_*conn*_ define the boundaries between the three regimes of *N*_*c*_(*ϵ*) (Fig. 1B): (i) for *ϵ* ≲ 12 *nm* the intra-clusters regime follows a Poissonian decay with 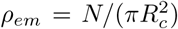 (see Supplementary Note 2); (ii) for intermediate values of *ϵ* the exponential super-clusters regime dominates and the fusion of connected clusters takes place; (iii) for *ϵ ≳* 60 *nm* we expect to observe the coalescence of super- and non-connected clusters in a second super-clusters regime; this should be captured again by a Poissonian decay for *p* = 0, and by an exponential for *p ≠* 0 (see Supplementary Note 3).

**FIG. 1:**
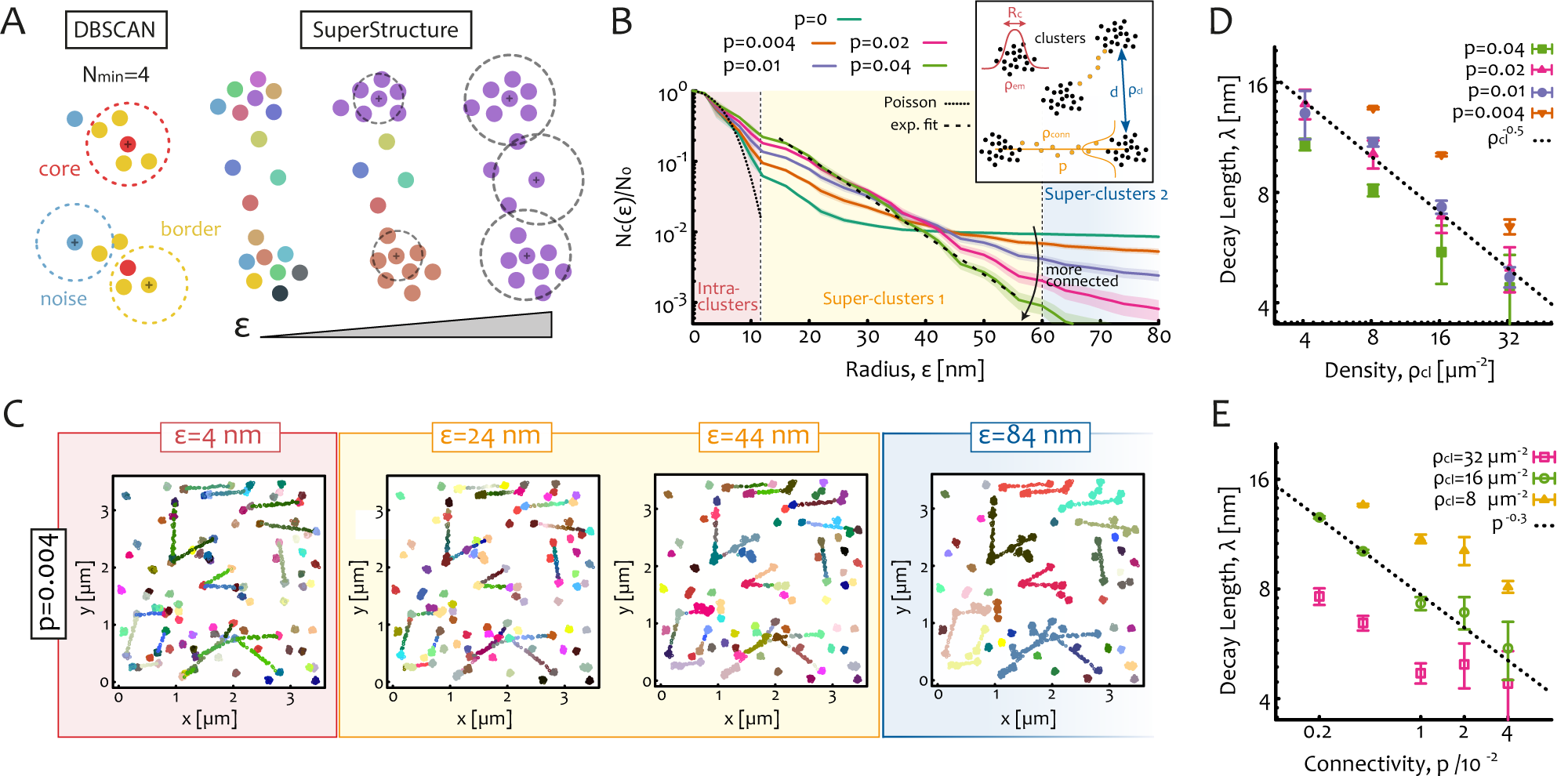
**A**. Left side: sketch of DBSCAN clusters detection for a choice of *ϵ* and *N*_min_ = 4; For every cluster DBSCAN identifies core (red) and border (yellow) points, as well as free noise (blue) points. Right side: sketch of SuperStructure algorithm for a pair of connected clusters. While *N*_*min*_ = 0 is kept fixed, the neighbour parameter *ϵ* is gradually increased. First points in clusters are merged, then clusters themselves are connected due to connection points. **B**. SuperStructure curves for random connected clusters for fixed average cluster radius *R*_*c*_ and inter-cluster distance *d* and different values of connectivity *p*. Three regimes can be distinguished: (i) intra-clusters (red), (ii) first super-clusters (yellow) and (iii) second super-clusters regime (blue). The decay in the intra-cluster regime corresponds to a Poisson Avoidance function with *ρ*_*em*_ = 16000 *µm*^*−*2^ (dotted line). The first super-clusters regime can be fitted by single exponentials (dashed line) which return an effective decay length *λ*. **C**. Snapshots for p=0.004 for *ϵ* = 4, 24, 44, 84 nm. **D**. *λ* versus *ρ*_*cl*_ has a 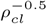 dependence for different *p*. **E**. *λ* versus *p* has a *p*^*−*0.3^a for different *ρ*.

Fig. 1B shows the exponential behaviour of the super-clusters regime (ii) for different values of *p*. Importantly, a larger connection probability *p* results in an effectively smaller decay length – or larger spatial rate of merging – for the super-clusters regime (ii). This strongly suggests that the effective decay length (or rate) mirrors the connectedness of the underlying super-structures (Fig. 1C). This decay length (plotted in Fig. 1D) results from the combined contribution of clusters density *ρ*_*cl*_ and connectivity *p*. A larger density of clusters can impact the decay length as much as a larger connectivity, as shown by simulations at fixed *p* and different *ρ*_*cl*_ (Fig. 1E and Supplementary Note 3). In particular, we find that the functional form of the decay length 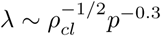 (Figs. 1D and E). In particular, the density contribution is 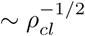as it depends on the typical distance between clusters and is relevant when comparing datasets with different cluster density (see below).

We then applied SuperStructure to dSTORM data acquired on three different nuclear proteins (Fig. 2A): the serine/arginine-rich splicing factor SC35, the heterogeneous nuclear RiboNuclear Protein hnRNP-C and hnRNP-U (also known as Scaffold Attachment Factor A, SAF-A). These proteins are abundantly expressed in the nucleus of human cells and are involved with RNA processing at different stages. SC35 is necessary for RNA splicing while hnRNPs are implicated in regulation and maturation of mRNA but also in chromatin structure [23, 32, 33]. In particular, SAF-A is thought to form a dynamic mesh that regulates large-scale chromatin organisation by keeping gene-rich loci in a decompacted state [23, 24]. Hence, understanding the organisation of this protein beyond the traditional single-cluster analysis is an important step towards understanding its function.

**FIG. 2:**
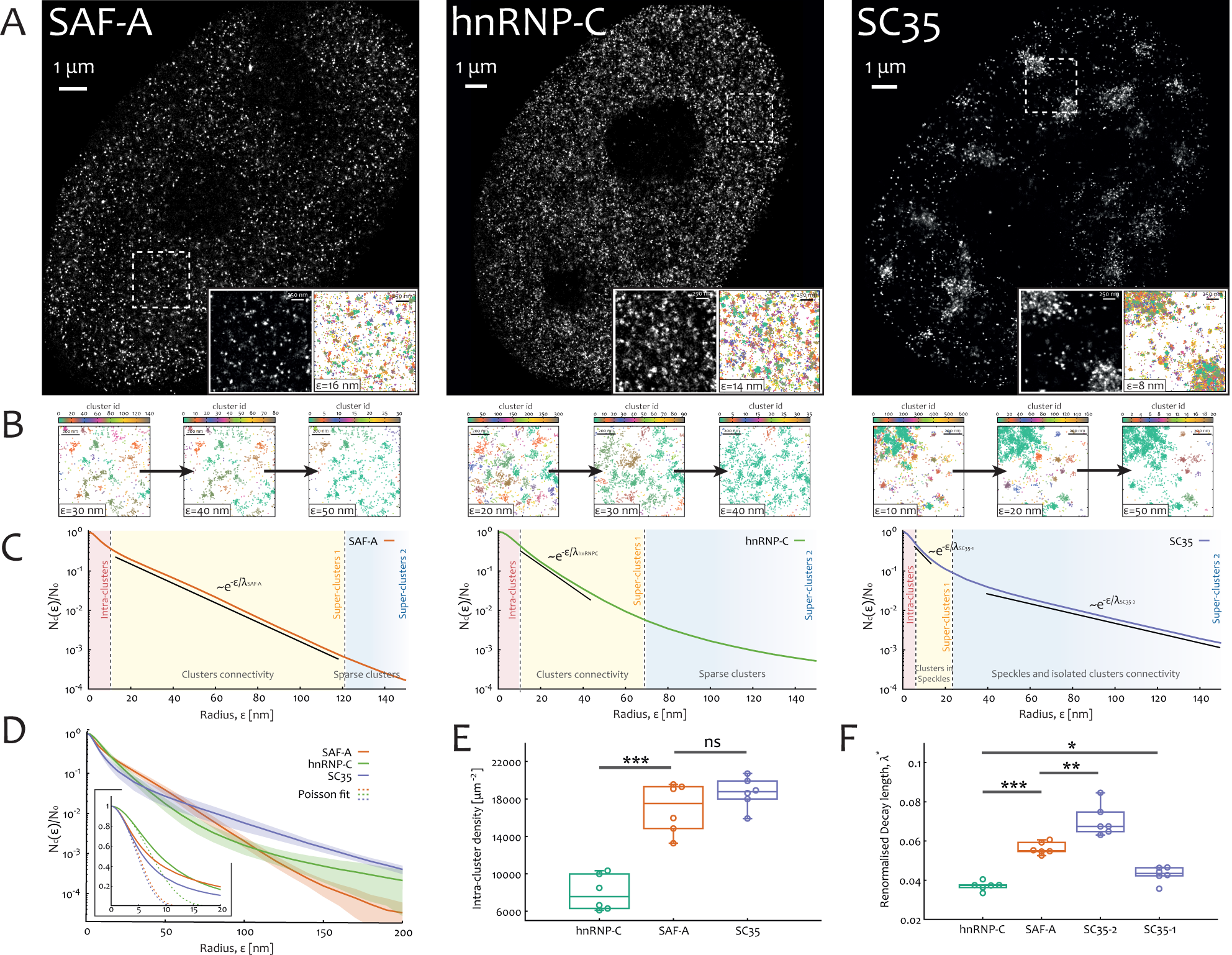
Application of SuperStructure algorithm to hnRNPc, SAF-A and SC35 super-resolution data. **A**. dSTORM reconstructed images by using a pixel size of 10 *nm*. Inset of 4 *µm*^2^ of super-resolution images and relative spatial positions of the data. Palettes represent the cluster id computed by running SuperStructure with *N*_*min*_ = 0 and a suitable value of *ϵ* for every protein (*ϵ* = 16 *µm* for SAF-A, *ϵ* = 14 *µm* for hnRNP-C, *ϵ* = 8 *µm* for SC35). **B**. Identified clusters for increasing values of *ϵ* in the regimes where clusters merge. **C**. Normalised average SuperStructure curves: the number of clusters normalised with the number of total points in the system while keeping *N*_*min*_ = 0 and increasing *ϵ* in the range [0 : 150] *nm*. The average is computed over 6 independent dSTORM acquisitions. Vertical dashed lines highlight different merging regimes: intra-clusters, first super-clusters and second super-clusters regimes. For SAF-A and hnRNP-C the exponential regime of clusters merging (first super-clusters) is highlighted. In case of SC35, two regimes are highlighted: the merging of clusters in speckles (first super-clusters) and the merging of speckles and isolated clusters (second super-clusters). **D**. Normalised average SuperStructure curves in the range [0 : 200] *nm*. Shaded region represents standard deviation from the average. Poisson fits of the intra-cluster regime at small *E* are shown in the inset. **E**. Intra-cluster density of emitters as calculate from Poisson fit at small *ϵ*. **F** Renormalised decay length (by cluster density) for the regimes highlighted in **C**. Note that significance is annotated as follows: *p*-*value ≤* 0.001 as *\* * \**, 0.001 *≤ p*-*value ≤* 0.01 as *\*\**, 0.01 *≤ p*-*value ≤* 0.05 as *\** and *p*-*value ≥* 0.05 as *ns*.

Average SuperStructure curves for these proteins are shown in Fig. 2C, where we highlighted the regimes discussed above. Both hnRNPs display a first super-clusters regime for which the curves decay as exponentials, suggesting that within this range distinct clusters tend to coalesce. Interestingly, SC35 displays exponentials with different characteristic rates in two distinct super-clusters regimes: one for intermediate *ϵ*, when clusters inside speckles merge (first super-clusters regime), and another one for high *ϵ* indicating that speckles merge together and with single clusters (second super-clusters regime). These connectivity properties were further confirmed by directly looking at the arrangement of identified clusters (inset of Fig. 2A and Fig. 2B).

We first fitted the intra-cluster regime with the Poisson function (see Supplementary Note 3 and 4). Interestingly, SAF-A and SC-35 form clusters with similar densities, while those of hnRNP-C are less dense (see Fig. 2D and E). Then, in order to have a quantitative description of the clusters/speckles connectivities, we calculated the decay length in the exponential regimes. However, a direct comparison is possible only by renormalising decay lengths by the cluster/speckles density (see also Supplementary Note 4). Fig. 2F highlights that while hnRNP-C has a short renormalised decay length due to the highly connected clusters, SAF-A displays a slower decay (larger *λ*^*\**^) due to a more weakly connected mesh. Finally, SC35 displays one (intra-speckles) very20connected and fast regime of the order of that of hnRNPs (small *λ*^*\**^) followed by a regime (inter-speckles) that is much slower compared to hnRNPs. In other words, our analysis revealed that the super-structures inside nuclear speckles are not more connected than those formed by hnRNPs but they are denser, as explained in Supplementary Note 4. Further, this method is also sensitive enough to distinguish the connectivity of two closely related wild-type hnRNPs in live cells.

SuperStructure was also evaluated on published dSTORM datasets on ceramids [34] (Supplementary Note 5). In this case we were able to confirm the quantitative results of the authors, but moreover we found a notable absence of connections between clusters, further supporting that the connections detected in hnRNP-U/C and SC35 are significant and potentially functional.

In summary, we have developed a novel analysis method that yields the degree of connectivity between clusters in SMLM data. Because parameter-free and based on a well established and highly popular clustering software (DBSCAN), we believe that our method will find broad use as standardised tool in SMLM data analysis and to tackle challenging datasets. We stress that the proposed method is ideal to quantify and compare different datasets without a priori assumptions. Additionally, we predict this method could be also applied for technical purpose, such as evaluation of fluorophores blinking quality.

## Methods

### SuperStructure algorithm and pipeline

The algorithm details and the complete pipeline are exhaustively discussed in Supplementary Note 1.

### Experimental details for generating experimental dSTORM dataset for SAF-A, hnRNP-C and SC-35

*Cells Preparation for dSTORM imaging*. RPE1 cells were grown overnight in a 8-well Nunc Lab-Tek II Chambered Coverglass (1.5 borosilicate glass) at 37 degrees at initial concentration of 10^5^ *cells/ml*. We fixed the cells for 10 minutes in 4% PFA, followed by washing in PBS, permeabilisation with 0.2% Triton-X for 10 minutes, washed in PBS again and blocked with 1% BSA for 10 minutes.

Immuno-fluorescence labelling was done by exposing the cells for 2 hours to (i) hnRNP-U polyclonal rabbit antibody A300-690A from *Bethyl Laboratories* at 10 *µg/ml* or (ii) hnRNP-C1/C2 (4F4) mouse monoclonal antibody sc-32308 from *Santa Cruz Biotechnology* at 0.2 *µg/ml* or (iii) SC-35 mouse monoclonal antibody ab11826 from *abcam* at 2 *µg/ml* and then washed. Then, cells were exposed for 1 hour to secondary antibody. The secondary antibody was made by anti-rabbit or anti-mouse F(ab) fragments fused to organic fluorophore CF647 at a stechiometric ratio of about 1.

Oxygen scavenger imaging buffer for dSTORM was prepared fresh on the day and the recipe employed was similar to that of Ref. [35]. We mixed (i) 5.3 *ml* of 200 mM Tris and 50 mM NaCl solution with (ii) 2 *ml* of 40% glucose solution, (iii) 200 *µl* of GLOX, (iv) 1.32 *ml* of 1M *β*-mercaptoethanol and (v) 100 *µl* of 50 *µg/ml* DAPI solution. The GLOX solution was made by mixing 160 *µl* of 200 mM Tris and 50mM NaCl with 40 *µl* of bovine liver catalase and 18 mg of glucose oxidase. The 8.8 *ml* final solution was suitable to fill the chambers of the 8-well dish and a cover glass was sealed at the top of the dish to prevent inflow of oxygen.

*dSTORM Acquisition*. We performed 3D STORM acquisitions using a Nikon N-STORM system with Eclipse Ti-E inverted microscope with laser TIRFilluminator (Nikon UK Ltd, Kingston Upon Thames, UK). We equipped the microscope with a CFI SR HP Apo TIRF 100x objective lens and applied a 1.5X additional optical zoom. The Z position was acquired by using a cylindrical astigmatic lens. Laser light was provided via a Nikon LU-NV laser bed with 405, 488, 561, 640nm laser lines. In particular, fluorophores were stochastically excited using the 640 nm laser beam. Images were acquired with an Andor iXon 897 EMCCD camera (Andor technologies, Belfast UK). The Z position was stabilised during the entire acquisition by the integrated perfect focus system (PFS).

For every nucleus, we acquired a stack of 20000 (256 pixels x 256 pixels) frames with 19 *ms* exposure time. Acquired images had a pixel resolution of 106 nm/pixel. For every condition (SAF-A, hnRNP-C, SC35 antibody labelling) we acquired 6 nuclei, i.e. 6 independent datasets.

#### Raw images and post-processing analysis

The raw stack of frames was analysed with the FIJI Thunderstorm plugin [36]. Frames were initially filtered by using Wavelet functions to separate signal from noise. The B-Spline order was set to 3 and the B-Spline scale to 2.0 as suggested in Ref. [36]. In order to localise the emitters centroids, we thresholded filtered images (threshold values was set 1.2 times the standard deviation of the 1st Wavelet function) and calculated the local maximum relative to the 8 nearest neighbours. Finally, we fitted the emitters signal distribution with elliptical gaussians (ellipses are necessary for z-position localisation) using the weighted least square method and by setting 3 *pixels* as initial fitting radius and 1.6 *pixels* as initial sigma.

Localised data was then post-processed using the same plugin. (i) We corrected the XY drift using a pair correlation analysis, (ii) filtered data with a position uncertainty *<* 40, (iii) restricted the z-position to the interval [*−*100 : 100] *nm* and projected the data in a 2-dimensional plane, as the z-axis precision is around 100 *nm*.

Reconstructed images shown in the main text were created by using the average shifted histograms method with a 10X magnification (10.6 nm/pixel).

## Acknowledgements

MM is a cross-disciplinary post-doctoral fellow supported by funding from the University of Edinburgh and Medical Research Council (core grant to the MRC Institute of Genetics and Molecular Medicine). SvdL is supported by the Academy of Medical Sciences/the British Heart Foundation/the Government Department of Business, Energy and Industrial Strategy/the Wellcome Trust Springboard Award (SBF003 \1163). N.G. is funded by the UK Medical Research Council (MC UU 00007/13). DM is supported by the Leverhulme Trust (Early Career Fellowship ECF-2019-088). The authors thank the support of the Scottish University Life Science Alliance through a technology seed grant Worktribe Project ID 8824507. The authors thank the ESRIC Imaging Team (IGMM section), in particular Matthew Pearson and Ann Wheeler for their support. The authors are grateful to Markus Sauer for providing with the ceramides data. MM and DM would also like to thank Ibrahim Cissé for an igniting discussion. The authors also thank discussions with Davide Marenduzzo’s group.

## Data Availability

The simulated and experimental datasets that support the findings of this study are available from the corresponding authors upon request.

## Author Contribution

M.M., D.M. and N.G. conceived the project. M.M. and D.M. analysed both simulated and experimental datasets. M.M., S.v.d.L. and D.M. generated the simulated dataset. M.M., E.L. and D.M. performed super-resolution experiments and localisation analysis. M.M., D.M., S.v.d.L. and N.G. wrote the manuscript with input from all the authors.

## Code Availability

The code for the generation of SuperStructure curves is available from *https://git.ecdf.ed.ac.uk/dmichiel/superstructure*.

## Additional Information

**Supplementary Notes** with the algorithm pipeline and the Super Structure analysis of the simulated/experimental datasets are available.

